# Blood variation implicates respiratory limits on elevational ranges of Andean birds

**DOI:** 10.1101/2021.09.30.462673

**Authors:** Ethan B. Linck, Jessie L. Williamson, Emil Bautista, Elizabeth J. Beckman, Phred M. Benham, Shane G. DuBay, L. Monica Flores, Chauncey R. Gadek, Andrew B. Johnson, Matthew R. Jones, Jano Núñez-Zapata, Alessandra Quiñonez, C. Jonathan Schmitt, Dora Susanibar, Tiravanti C. Jorge, Karen Verde-Guerra, Natalie A. Wright, Thomas Valqui, Jay F. Storz, Christopher C. Witt

## Abstract

The extent to which species ranges reflect intrinsic physiological tolerances is a major, unsolved question in evolutionary ecology. To date, consensus has been hindered by the limited tractability of experimental approaches across most of the tree of life. Here, we apply a macrophysiological approach to understand how hematological traits related to oxygen transport shape elevational ranges in a tropical biodiversity hotspot. Along Andean elevational gradients, we measured traits that affect blood oxygen-carrying capacity—total and cellular hemoglobin concentration and hematocrit—for 2,355 individuals of 136 bird species. We used these data to evaluate the influence of hematological traits on elevational ranges. First, we asked whether hematological plasticity is predictive of elevational range breadth. Second, we asked whether variance in hematological traits changed as a function of distance from the midpoint of the elevational range. We found that the correlation between hematological plasticity and elevational range breadth was slightly positive, consistent with a facilitative role for plasticity in elevational range expansion. We further found reduced local variation in hematological traits near elevational range limits and at high elevations, patterns consistent with intensified natural selection, reduced effective population size, or compensatory changes in other cardiohematological traits with increasing distance from species-specific optima for oxygen availability. Our findings suggest that constraints on hematological plasticity and local genetic adaptation to oxygen availability promote the evolution of the narrow elevational ranges that underpin tropical montane biodiversity.

## Introduction

When, if ever, are species ranges limited by intrinsic physiological tolerances? Correlative niche models have demonstrated the pervasive influence of climate on plant and animal distributions (reviewed in Elith & Leathwick, 2009), but the specific mechanisms that link abiotic variables to organismal fitness and spatial variation in population viability remain elusive for most taxa (Bozinovic et al., 2011; Bozinovic & Naya 2015). The challenge is acute because the vast majority of species are impractical for experimental study. In these cases, macrophysiological data—measurements of physiological trait variation at large spatial and phylogenetic scales—can provide valuable insights on the functional underpinnings of biogeographic patterns (Chown et al., 2004; Gaston et al., 2009).

Few biogeographic patterns are as striking as pervasive elevational specialization in tropical vertebrates. In the Peruvian Andes, for example, birds have a median elevational range breadth of approximately 1100 meters, despite over 5000 meters of habitable elevation gradient below the line of permanent snow and ice (Stotz et al., 1996). This specialization contributes to extraordinary beta diversity: along a single surveyed transect near Manu, Peru, 66 hummingbird species occur, but no more than 30 at a single elevation (Walker et al., 2006). Because rapid species turnover is correlated with a dramatic environmental gradient (McNew et al., 2021), biologists have long hypothesized that narrow breadth of tolerance to one or more abiotic variables contributes to range limits (von Humboldt, 1838; Janzen, 1967; Terborgh, 1971; Jankowski et al., 2013). Yet to date, research on the proximate causes of range limits has largely focused on biotic interactions (Terborgh & Weske, 1975; Freeman et al., 2016), complemented by a handful of studies on metabolic and thermal physiology with largely equivocal results (Londoño et al., 2015; Londoño et al., 2017; Wolf et al., 2020; Gutierrez-Pinto et al., 2021).

One environmental variable with well-characterized consequences for organismal physiology and an obvious influence on species distributions at coarse spatial scales is the partial pressure of oxygen (*P*O_2_), which systematically declines with increasing elevation. Lowland organisms show a variety of plastic acclimatization responses to low *P*O_2_, including interrelated changes in numerous respiratory, cardiovascular, and metabolic traits (Storz et al., 2010; Storz & Scott, 2019). These plastic responses are often maladaptive, such as when excess erythropoiesis leads to harmful increases in blood viscosity (Storz et al., 2010). On longer time scales, evolutionary shifts in elevation are linked with predictable patterns of adaptation to the hemoglobin protein that shift oxygen-binding affinity to improve oxygenation, in birds if not mammals (Projecto-Garcia et al., 2013; Natarajan et al., 2016; Storz, 2016; Storz, 2018). The fitness tradeoffs associated with physiological adaptation to a particular *P*O_2_ regime could promote the evolution of elevational specialization in vertebrates. If so, blood phenotypic variation across the elevational range of species may bear the signature of this evolutionary process. For example, systematic variation in individual plasticity in hemoglobin concentration across species is a trait that could affect overall acclimatization capacity and, by extension, environmental niche breadth. Similarly, the relationship between local (elevation-specific) variation in hemoglobin concentration and relative position within a species’ elevational range could indicate stronger natural selection near range limits (Hoffman & Blows, 1994; Pennington et al., 2021). These hypotheses await validation.

Here, we apply a macrophysiological approach to understand how blood oxygen-carrying capacity might shape geographic distributions in a biodiversity hotspot. We measured total and cellular hemoglobin concentration and hematocrit variation for 2,355 individuals of 136 bird species and used these data to evaluate the influence of hematological traits on elevational ranges. First, we asked whether hematological plasticity is predictive of elevational range breadth, using the rate of increase of hematological trait values with elevation as a proxy for the species-wide mean of individual plasticity measurements. Because a hypoxia-induced increase in [Hb] can contribute to hypoxia acclimatization by increasing arterial O_2_ content—augmenting convective O_2_ transport when paired with other cardiovascular adjustments (Stembridge et al. 2019; Gonzalez et al. 2002; Tate et al. 2020)—plasticity in [Hb] could be positively correlated with elevational range breadth. Alternatively, [Hb] might instead be a hypoxia-responsive trait that reflects overall levels of tissue O_2_ delivery. In this scenario, species with O_2_- transfer pathways that are robust to hypoxia would show little change in [Hb] with increasing elevation and occupy a broader range of *P*O_2_ regimes, while those with limited hypoxia tolerance would show a higher rate of increase in [Hb] as a function of elevation. Plasticity in [Hb] would thus be negatively correlated with elevational range breadth.

Second, we asked whether variance in hematological traits changed as a function of distance from the midpoint of the elevational range. Several non-mutually exclusive mechanisms could lead to nonrandom patterns of variance in trait values across elevation. Under the reasonable assumption that these values directly reflect underlying additive genetic effects, variance could be constrained at range margins by local reductions in effective population size due to source-sink dynamics or other demographic processes that may or may not be related to diminished respiratory performance (Hoffman & Blows, 1994; Gaston, 2003; Vucetich & Waite, 2003). Alternatively, populations near elevational range limits might experience stronger natural selection to optimize blood oxygen-carrying capacity than those near elevational range centers (Hoffman & Blows, 1994; Pennington et al., *in press*). If we assume trait values have no genetic basis, decreased variation may still reflect an increase in the relative importance of compensatory changes in other cardiohematological traits. Agnostic to mechanism, we predicted that departures from species-specific optima for respiratory performance would lead to a negative correlation between distance from range center and local variation in hematological trait phenotypes.

## Methods

### Haematological measurements

From 2006 to 2020, we measured total blood hemoglobin concentration ([Hb]) and the percentage of hematocrit (Hct) for birds captured during collaborative fieldwork in Peru by the Museum of Southwestern Biology (MSB) in New Mexico, USA, and the Centro de Ornitolog*í*a y Biodiversidad (CORBIDI) in Lima, Peru. We conducted this work with assistance from numerous researchers associated with these institutions—whose contributions we gratefully acknowledge here and describe in detail elsewhere (Witt et al., *in prep*.)—and under the following permits from Peru’s management authorities: 004-2007-INRENA-IFFS-DCB, 135-2009-AG-DGFFS-DGEFFS, 0377-2010-AG-DGFFS-DGEFFS, 0199-2012-AG-DGFFS-DGEFFS, 006-2013-MINAGRI-DGFFS/DGEFFS, 280-2014-MINAGRI-DGFFS-DGEFFS, and 2022-RDG N° 405-2017-SERFOR-DGGSPFFS.

As quickly as possible after capture, we punctured the brachial vein on the underside of each bird’s wing and collected whole blood using heparinized microcapillary tubes (for Hct) or spectrophotometer cuvettes (for [Hb]). We centrifuged microcapillary tubes for 5 minutes at 13,000 rpm to separate red blood cells from plasma and used digital calipers to quantify the relative volume of each, averaging two samples per bird to arrive at our final estimate of Hct as a percentage. We measured [Hb] (g/dL blood) for a ∼5 μL blood sample with the Hemocue HB201 analyzer and associated hemoglobin photometer. As this proprietary method generates values approximately 1.0 g/dL higher than measures from a direct cyanomethaemoglobin spectrophotometer (Simmons & Lill, 2006), we corrected the resulting estimates by subtracting this quantity (DuBay & Witt, 2014). Specimens and tissues are housed at the MSB and CORBIDI and specimen data are detailed in a separate data publication (Witt et al., *in prep*.)

### Data filtering and preparation

To ensure only high-quality measurements were included in our analyses, we applied a series of filters to the full dataset of hematological traits. We first removed all records that lacked a measurement for either [Hb] or Hct, and then used these values to calculate MCHC (mean cellular hemoglobin concentration; 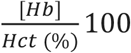 (Campbell & Ellis, 2007). We then removed outliers using a set of thresholds for the minimum and maximum allowable value of each hematological trait, which were derived by visualizing distributions and Q-Q plots and assuming significant deviations from normality represented measurement error, unhealthy individuals, or otherwise unusable data. These thresholds were: [Hb] ≤ 11 or ≥ 24; Hct ≤ 0.3 or ≥ 0.8; and MCHC ≤ 22 or ≥ 42. After applying outlier filters, we dropped any species with < 5 remaining records and merged the resulting dataset with estimates of elevational range breadth compiled from the literature and verified by expert opinion (Stotz et al., 1996; Schulenberg et al., 2010), expanding these estimates as necessary to encompass our own observations when they were fell outside the bounds of previously published datasets.

Because we were interested in using rate of change in each hematological trait per unit elevation as a proxy for hematological plasticity, and therefore wished to avoid biasing estimates through the influence of a single erroneous measurement, we next ran simple linear regressions of [Hb], Hct, and MCHC against the elevation they were recorded at and removed all points with a Cook’s distance of 4/*n* (where *n* is sample size). For individuals in our dataset that had a bursa of Fabricius—suggesting they were immature, and potentially exhibiting anomalous hematological trait values (Fair et al., 2007)—we used a more conservative Cook’s distance cutoff of 3.5/*n*. We then regressed [Hb], Hct, and MCHC against elevation a second time, and generated a new dataset where each row referenced a single species, with estimates of the slope of each hematological trait and its associated standard error. After dropping species lacking data from at least two elevations >200 meters apart—a span chosen to reflect evidence of fine-scale physiological sensitivity to *P*O_2_ in vertebrates (Gassmann et al., 2019)—we combined this dataset of hematological plasticity and elevational range breadth values with estimates of the median elevation of each species’ range (calculated as *Elev*_*min*_ + *Elev*_*breadth*_/2) and its average mass (calculated as the arithmetic mean of the recorded mass of all individual birds in our final filtered dataset). Lastly, we calculated the proportion of each species’ total elevational range represented by individuals in our dataset, a metric we included to evaluate the possible effects of sampling bias on our data.

To understand variation in hematological traits within species at a given approximate elevation, we generated a second dataset using the above outlier filters and minimum per-species sample size. We then divided the sampled elevational range of each species into 100-meter bins, discarding the remainder in the event of a non-integer quotient. In each bin with a minimum of 6 records we calculated the coefficient of variation (hereafter “CoV”) as 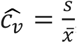 where *s* is the standard deviation of the sample and 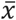 is the arithmetic mean. This resulted in a dataset where each row referenced a bin-specific CoV value, which we associated with the mean elevation of all individuals sampled within that 100-meter bin, their species, and the relative distance of the mean elevation of each bin from the nearest elevational range limit.

### Bayesian models

To evaluate the hypothesis that hematological plasticity was predictably related to elevational range characteristics, we built a set of generalized multivariate linear models in the R package brms v.2.13.5 (Bürkner, 2017), which is a wrapper for the statistical programing language Stan. For each hematological trait ([Hb], Hct, and MCHC), we modeled elevational plasticity (*P*) using a Student’s t-distribution for outcome variables and regularizing priors:

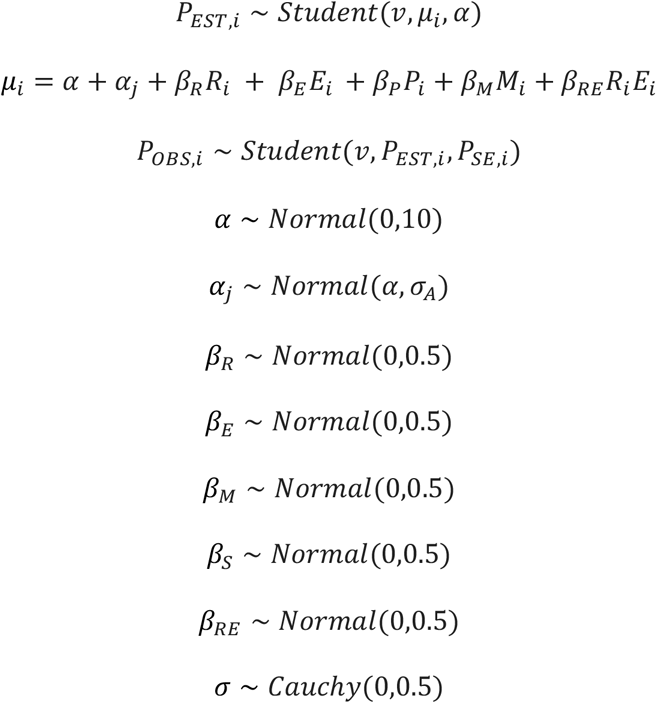

Here, the predictor *R* is elevational range breadth, *E* is median range elevation, *S* is proportion of elevational range sampled, *M*is mass, and *A* is a covariance matrix of phylogenetic distance among taxa. We generated this matrix by applying the function maxCladeCred() from the R package phangorn v.2.5.5 to 10,000 trees from the Hackett “backbone” phylogeny downloaded from BirdTree.org, and pruned it to the subset of species present in our dataset using the keep.tip() function in the R package ape v.5.3 (Paradis et al., 2004; Schliep, 2011; Jetz et al., 2012). Because this proxy for hematological plasticity is associated with substantial uncertainty, we model measurement error by treating the outcome variable as a vector of parameters with the likelihood *P*_*EST,i*_ ∼ *Student*(*v, μ, α*), which is given a prior that treats our observed data as drawn from a Student’s t-distribution with unknown mean *P*_*EST,i*_, and the calculated standard error of the regression (*P*_*SE,i*,_). In this way, uncertainty in the outcome variables is incorporated into regression parameters but is itself influenced by the linear model (McElreath, 2020).

This “full” model (1) reflects the more specific hypothesis that all predictors and phylogeny influence *P*. To evaluate whether predictive power was improved by simplifying model structure, we built reduced models that (2) included the interaction term between *R* and *E* but did not include phylogenetically correlated intercepts; (3) did not include either the interaction term between *R* and *E* or phylogenetically correlated intercepts; and (4) included phylogenetically correlated intercepts but did not include the interaction term between *R* and *E*. We then created a null model (5) where outcome variables were solely influenced by phylogenetically correlated intercepts. As these five model backbones were repeated for each of the three outcome variables, we fit a total of 15 plasticity models.

We built similar models to evaluate whether variance in hematological traits changed as a function of distance from the midpoint of the elevational range. We modeled the CoV (*V*) in [Hb], Hct, and MCHC within a given elevational band using a logarithmic distribution and without modeling measurement error:

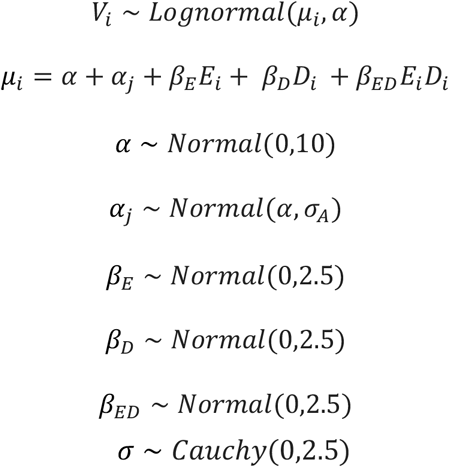

In the full model (1) above, predictor *E* is mean elevation of samples in a given bin, *D*is the mean distance of samples from the nearest elevational range limit, and the intercept terms are defined as previously described. We created three reduced models by (2) excluding phylogenetically correlated intercepts, (3) excluding the interaction term between *E* and *D*, and (4) excluding both phylogenetically correlated intercepts and interaction term between *E* and *D*. We compared all four models with predictors to a null model (5) that included a phylogenetically correlated intercept alone. As with plasticity models, this created a total of 15 additional models, with five models each for the outcome variables [Hb], Hct, and MCHC.

Prior to fitting brms models, we standardized each predictor, and scaled outcome variables to the same order of magnitude. We fit all models using two MCMC chains with 5000 generations of warmup and 5000 generations of sampling each. We evaluated convergence and stationarity by examining trace plots, ensuring effective sample sizes were sufficiently high (>1000), and verifying for each model that the Gelman-Rubin diagnostic 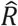 was less than or equal to 1. We further evaluated model fit by performing a posterior predictive check of the distribution of each outcome variable and screened for sampling issues by creating scatterplots of the relationships among variables from MCMC draws.

To compare the predictive ability of alternate plasticity and CoV models for the same outcome variables, we applied leave-one-out cross validation information criteria (LOOIC) and assessed differences in the expected log pointwise predictive density (ELPD) using the function loo() in brms. For the full model for each outcome variable and summary statistic, we used the function median_hdi() in the R package tidybayes v.2.1.1 to calculate the 95%, 80%, and 50% credible intervals of the posterior probability distribution of each predictor and interaction. We visualized these distributions using the tidybayes and ggplot v.3.3.2.

For each model with a predictor whose 80% confidence interval did not overlap zero, we used the predict() and fitted() functions in brms to generate fit curves with probability distributions, with and without incorporating residuals, respectively. To do so, we used parameter estimates from the best-fit model (**Table 1**) to predict new observations of a given outcome variable across 3 standard deviations of the credible predictor while holding all other predictors constant at their mean value. For interaction terms that were credible at the 80% threshold, we repeated this approach three times for a single predictor in the interaction, holding the remaining predictor at −2 standard deviations, its mean value, and +2 standard deviations. In this way, we were able to visualize the effect of one predictor on the change in the slope of the relationship between a second predictor and its outcome variable.

**Table 1:** Relative performance of models for 1) hematological trait plasticity to elevation (indicated by the prefix Δ) and 2) local variation in hematological traits at a given elevation (indicated by the prefix CoV). Values represent the change in expected log pointwise predicted density (ELPD) from leave-one-out cross-validation of Bayesian models, with associated standard errors reported in parentheses. Variable-specific predictors and interaction terms are described in Methods. ELPD values reflect the difference between each model for a given response variable and the best fit model among all comparisons (highlighted here in bold).

| Model Description |  |  |  | Response |  |  |  |  |  |
| --- | --- | --- | --- | --- | --- | --- | --- | --- | --- |
| Number | Predictors | Interaction | Phylogeny | $\Delta$ [Hb] | $\Delta$ Hct | $\Delta$ MCHC | CoV [Hb] | CoV Hct | CoV MCHC |
| 1 | X | X | X | -1.8 (1.0) | -1.5 (0.8) | -2.9 (1.9) | -0.5 (0.8) | -1.2 (1.0) | -1.6 (0.6) |
| 2 | X | X |  | -1.1 (0.4) | -1.1 (2.1) | -2.2 (1.9) | **0** | -0.2 (0.9) | -0.7 (0.6) |
| 3 | X |  |  | **0** | -0.8 (2.1) | -1.8 (1.6) | -0.6 (1.3) | **0** | **0** |
| 4 | X |  | X | -1.0 (0.8) | -1.1 (0.8) | -2.6 (1.7) | -1.0 (1.5) | -0.1 (0.6) | -0.9 (0.4) |
| 5 |  |  | X | -1.7 (3.0) | **0** | **0** | -3.3 (3.0) | -6.4 (3.3) | -3.8 (3.2) |

## Results

### Data filtering

Our initial dataset included hematological trait measurements from 5,927 individual birds representing 656 unique species, with sampling from 107 unique localities ranging from 39 to 4578 meters above sea level (masl). After filtering, the reduced dataset used to model hematological plasticity to elevation included 2,355 hematological trait values from 136 species, collected from 39–4578 masl. Our reduced variance dataset included 118 CoV estimates for each hematological trait from 73 species. The minimum median elevation of the bins used to calculate CoV values was 250 meters (m), and the maximum median elevation was 4350 m.

### Plasticity and variance of hematological traits

In general, hematological traits were positively correlated with elevation within species following data filtering (**Figure 2; Figure 3**). The median value of the slope of [Hb] regressed against elevation (m) was 0.7 g/dL/1000 m, with a 50% interquartile range (IQR) of −0.4 to 1.7; for Hct, the median slope was 0.02% per 1000 m, with a 50% IQR of −0.01% to 0.05%. MCHC was less sensitive to elevation, with a mean slope of 0.07 g/dL/1000 m, and a 50% IQR of −0.00123 to 0.00121. The CoV was roughly comparable among all hematological traits, with a median value of 0.0752 for [Hb] (50% IQR: 0.0617 to 0.0983), 0.0743 for Hct (0.0579 to 0.0982), and 0.0497 for MCHC (0.0387 to 0.0665).

**Figure 1.**
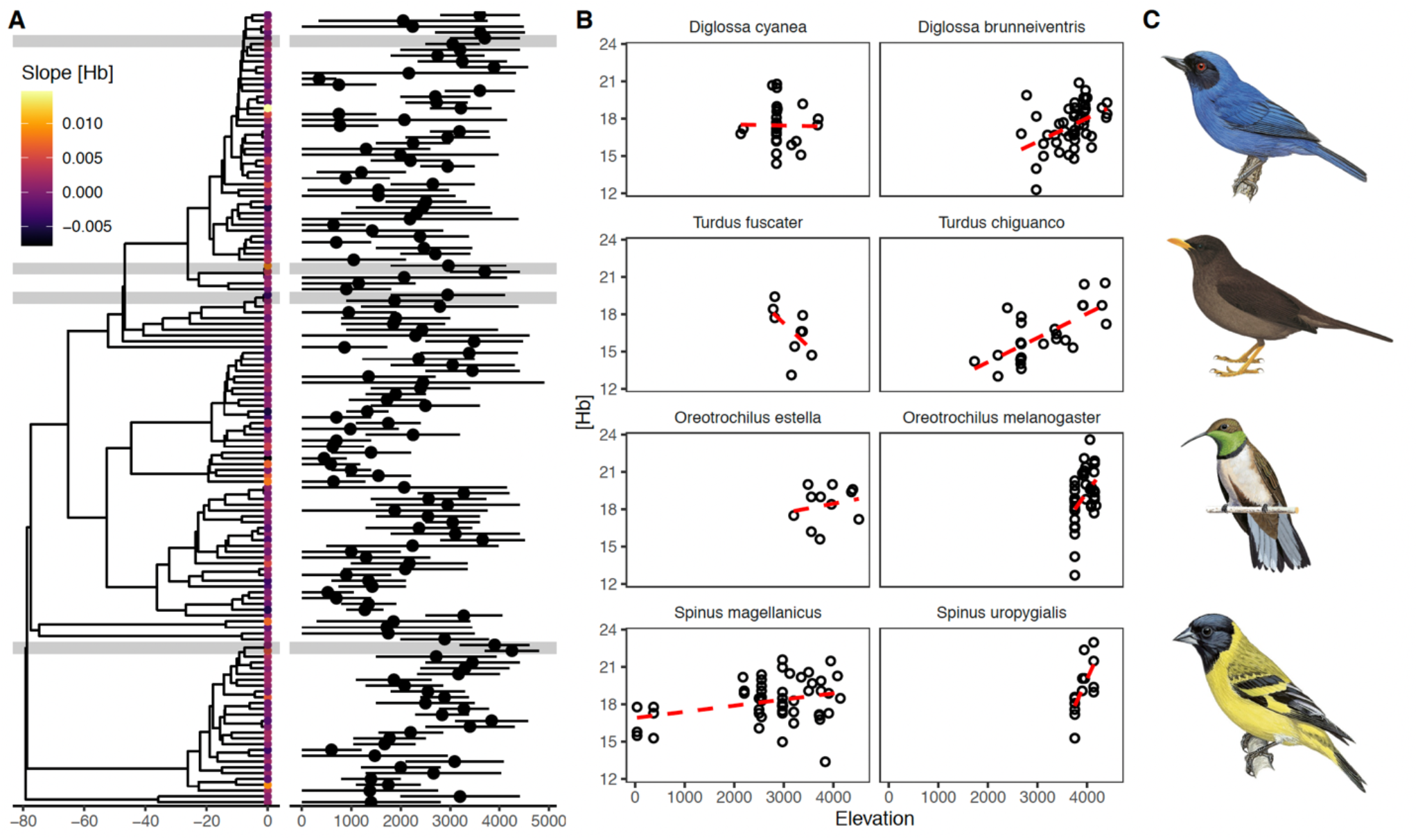
A) Low phylogenetic signal (λ=0.3) in plasticity of total blood hemoglobin concentration ([Hb]) to change in elevation across 137 species of Andean birds. Elevational range breadth and median range elevation (masl) are plotted adjacent to their corresponding species; values on the x-axis of the phylogeny represent millions of years ago before present. B) Examples of interspecific variation within genera in plasticity of [Hb] to elevation, with each genus’ position in the phylogeny highlighted in gray in corresponding vertical order to their respective plots. C) Illustrations of representative members of each genus from Birds of the World and reproduced courtesy of Cornell Lab of Ornithology (©).

**Figure 2.**
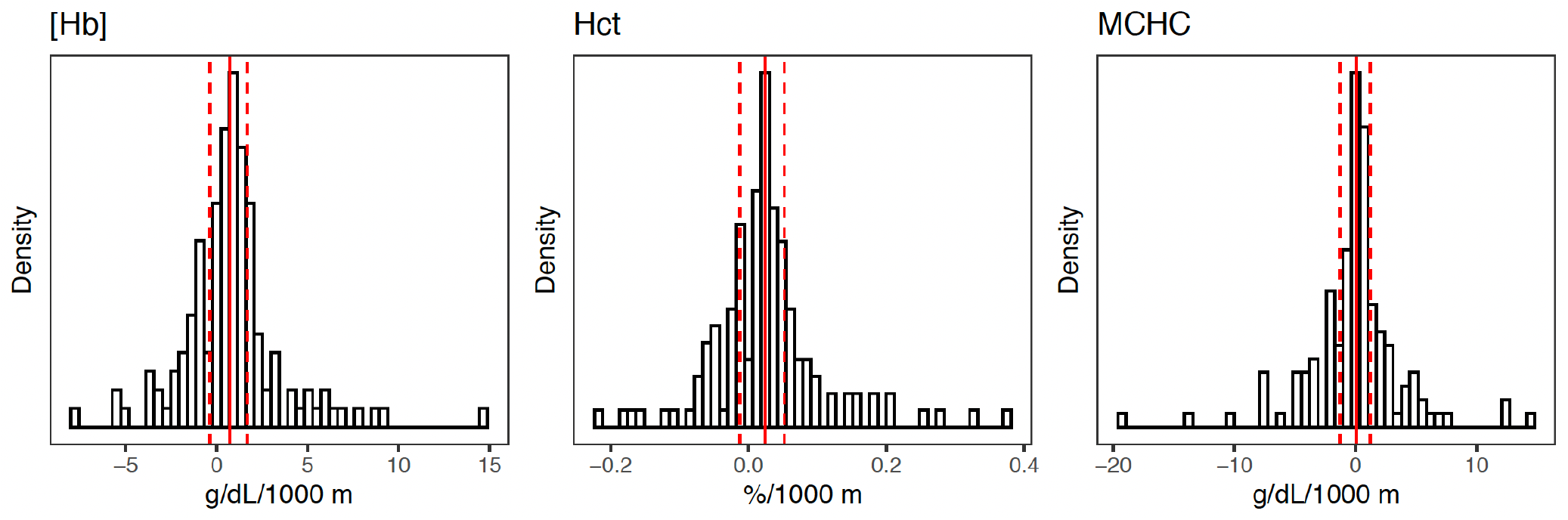
Observed distributions of hematological plasticity across elevation. Solid red lines indicate the median value and are flanked by dashed red lines that mark the 50% interquartile range (IQR).

**Figure 3.**
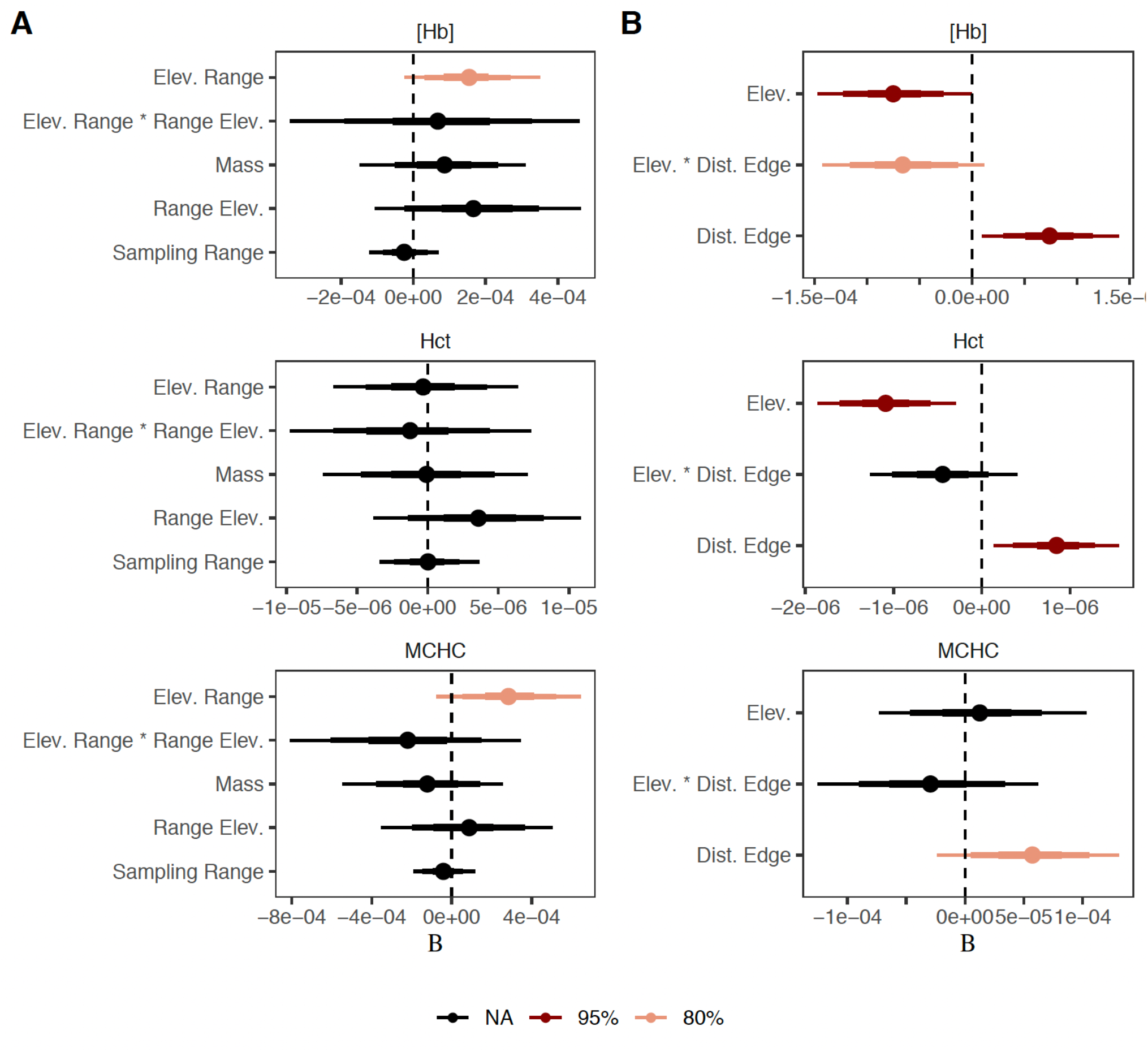
Posterior effect sizes for predictors in Bayesian models of A) hematological trait plasticity across elevation and B) the coefficient of variation in hematological traits within a given 100-meter elevational bin. Median estimates are indicated by points; decreasing line thickness away from the median indicate the 50%, 80%, and 95% credible intervals for each estimate. Predictors with 95% credible intervals that fall entirely above or below 0 are colored in dark red, while those with 80% credible intervals that fall entirely above or below 0 are colored in pink.

### Bayesian models

A model of [Hb] plasticity that included fixed effects for elevational range breadth, median range elevation, mass, and proportion of elevational range sampled—but not a term for the interaction between elevational range breadth and median range elevation, or phylogenetically correlated intercepts—had the highest ELPD, albeit with substantial standard error associated their improved performance over a null model (ΔELPD < 2*SE**; Table 1**). Elevational range breadth had a positive effect, credible at the 80% level. No predictors had a credible influence on Hct plasticity, which was best predicted by phylogenetically correlated intercepts alone, though again uncertainly (ΔELPD < 2*SE**; Table 1**). While a null model of MCHC plasticity also outperformed all others, elevational range breadth had a positive effect on MCHC plasticity with predictors, credible at the 80% level (**Figure 3**). As estimated in models that included all predictors and interaction terms, phylogenetic signal (lambda; λ) was 0.03 (95% credible interval=0.00-0.01) for [Hb], 0.19 for Hct (95% credible interval=0.02-0.43) for Hct, and 0.04 for MCHC (95% credible interval=0.00-0.12).

Models of the CoV of hematological traits that included fixed effects were universally better fits for our data than a phylogeny-only null model, though standard errors for differences in ELPD among models were large (all ΔELPD < 2*SE). The CoV of [Hb] within a given elevational band was best predicted by a model with fixed effects for elevation, distance from the nearest elevational range limit, and their interaction (**Table 1**). The mean elevation of samples had a negative influence on the CoV of a given 100-meter bin, with a 95% confidence interval that did not overlap with zero. Distance from the nearest elevational range limit had a positive influence on CoV, with a 95% confidence interval with both upper and lower bounds greater than zero (**Figure 3**). The interaction of elevation and distance from elevational range limit had a negative effect, with an interval that was credible at the 80% level, but not at the 95% level (**Figure 4**).

**Figure 4.**
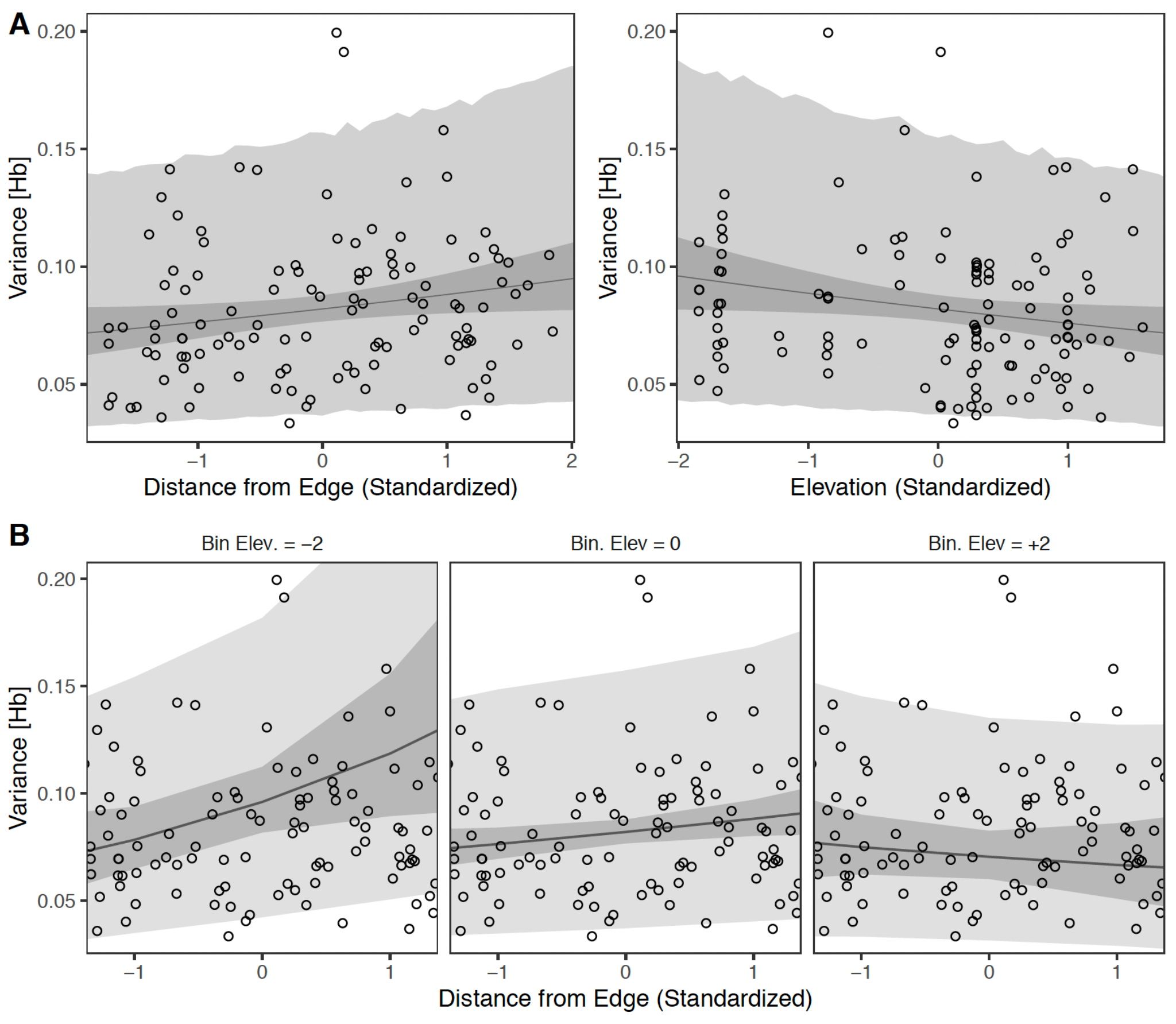
A) Predicted effects of distance from edge (left) and elevation (right) on variance in total hemoglobin concentration ([Hb]). The curves in each plot represent the contribution of the variable on the x-axis to the outcome variable while holding all other predictors constant at their mean value. B) The effect of elevation on the contribution of distance from edge to variance in [Hb]. Panes from left to right show that increasing the standardized value of elevation decreases the slope of the relationship between distance from edge and variance in [Hb].

The sign of the effect of these predictors was identical for the full model of CoV of Hct at a given elevation (**Figure 3**), though the model with the highest ELPD did not include an interaction term (**Table 1**). In the full model, elevation had a negative effect on the CoV of Hct, credible at the 95% level. Distance from the nearest elevational range limit was positively associated with CoV, and also credible at the 95% level; the effect of the interaction of elevation and distance from nearest elevational range limit was negative, with a 95% CI that did not overlap zero. Lastly, the best model for the CoV of MCHC included fixed effects (but not an interaction term or phylogenetically correlated intercepts; **Table 1**); distance from nearest elevational range limit had a positive effect, credible at the 80% level (**Figure 3**). The phylogenetic signal in models of the local CoV of hematological traits with all predictors and an interaction term was 0.01 (95% credible interval=0.00-0.03) for [Hb], 0.01 for Hct (95% CI=0.00-0.03) for Hct, and 0.01 for MCHC (95% CI=0.00-0.04).

## Discussion

The role of physiological tolerances in shaping species ranges remains poorly known for the vast majority of taxa, making it difficult to discern whether there are functional underpinnings for prominent biogeographic patterns. In a large dataset of intraspecific physiological measurements in Andean birds, we found hematological plasticity and local variation in hematological trait phenotypes were predictably associated with elevational range characteristics (**Figures 3-4**). Our study suggests hematological plasticity and genetic adaptation of hematological traits to a narrow range of oxygen partial pressure contributes to determining species elevational niches, linking a prominent dimension of the abiotic niche of montane vertebrates with realized geographic ranges.

Across 136 species, plasticity and elevational range breadth were positively correlated in [Hb] and MCHC, a pattern that supports the hypothesis that enhanced acclimatization ability may facilitate range expansion into novel *P*O_2_ regimes (**Figure 3a**). We consider this an intuitive, if tentative, result, as niche breadth is often correlated with the degree of plasticity in other contexts (Williamson & Witt, 2021). For example, an influential paper by Van Valen compared populations of six species of passerine birds co-distributed in island-mainland pairs, finding broad support for the hypothesis that functional morphological variation (as measured by the intraspecific coefficient of variation for a given trait) was “controlled to a significant extent by the adaptive diversity of the niche” (Van Valen 1965). While Van Valen assumed genetic control of the bill traits in his study, he acknowledged that under certain circumstances non-heritable plasticity might produce a similar pattern. In stickleback fishes, common garden experiments suggest phenotypic plasticity has evolved repeatedly in generalist populations (Svanbäck & Schluter, 2012), a finding both consistent with theory (Tienderen, 1997) and other empirical work (Bradshaw, 1965; Balaguer et al., 2001). In particular, plasticity appears to aid range expansion in many invasive species (Richards et al., 2006; Davidson et al., 2011; Knop & Reusser, 2012).

Alternatively, if the broad ranges of elevational generalists are ephemeral products of recent population expansions (Gadek et al., 2018), the positive correlation between elevational range breadth and hematological plasticity might be driven by temporal bias, as species with narrower, more stable ranges evolve plasticity-suppressing adaptations (i.e., through genetic compensation; Storz & Scott, 2020). We think this is unlikely: in species or lineages with long-term high-elevation ancestry, reduced hematological plasticity across elevation appears to instead reflect changes in other convective and diffusive steps in the O_2_-transport pathway that help sustain adequate levels of tissue oxygenation in spite of environmental hypoxia, dampening the hypoxic stimulus to increase [Hb] (either via increased erythropoiesis, contracted plasma volume, or both; Stembridge et al., 2019). Furthermore, we would expect species with high median range elevations to show reduced hematological plasticity if genetic compensation were an important predictor of trait values. This appears not to be the case (**Figure 3a**).

We find the result of reduced local variation in hematological traits near elevational range limits and at high absolute elevation (**Figure 3b**) similarly intuitive but more difficult to interpret, as competing processes could either exclusively or cooperatively generate a comparable pattern. As hematological traits are highly labile and responsive to changes in tissue *P*O_2_, decreasing variance with increasing proximity to elevational range limits might primarily reflect an increased role for compensatory changes in other cardiohematological traits (such ventilation and pulmonary O_2_ diffusion) that affect the hypoxic stimulus for activating erythropoiesis or changes in plasma volume. Assuming trait values at least partly reflect additive genetic variation, lower CoV values might also result from neutral demographic processes that lead to genome-wide reductions in effective population size (*N*_e_). Low *N*_e_ is expected near range limits under the predictions of extensions of the central-marginal hypothesis and its variants (Hengeveld & Haeck, 1982; Brown, 1984; Hoffman & Blows, 1994; Gaston, 2003; Vucetich & Waite, 2003), and while this pattern appears far from universal (Eckert et al., 2008)—especially across elevational gradients (Freeman & Beehler, 2018)—it remains a viable alternative explanation.

While acknowledging these possibilities, the scenario we consider most intriguing is that lower CoV values at range limits are a signature of spatially varying strength of selection on respiratory phenotypes, implicating hematological traits in the maintenance of elevational distributions in Andean birds (**Figure 3b, Figure 4a**). Specifically, reduced variation in [Hb] and Hct near elevational range limits might suggest an increasing fitness cost for phenotype-environment mismatches and, by extension, suggests that species’ distributions may be partially limited by a failure to adapt or acclimate to *P*O_2_ conditions beyond their current elevational niche. Likewise, reduced variation in [Hb] and Hct at high absolute elevation (**Figure 4b**) would be expected if the nonlinear decline in arterial O_2_ saturation with increasing elevation leads to stronger selection regardless of relative range position. Elevational variation in hematological trait phenotypes may also be influenced by locally adaptive genetic variation along the elevational gradient (Schweizer et al. 2019; Lim et al. 2021). If so, the higher CoV values at elevational range center in our models could partly reflect increased heterozygosity of causal loci that are subject to variable selection across elevation, or a mosaic of alleles that are favored at upper and lower elevations, respectively.

More broadly, the spatial distribution of ecologically meaningful trait values and functional genetic variation is a potentially rich source of information on range limiting mechanisms. Though studies of range limiting mechanisms remain rare outside of a handful of well-studied taxa, a recent metanalysis by Pennington and colleagues (*in press*) found quantitative genetic variation declined from geographic range centers to range margins but increased towards niche limits. Under the simplifying assumption that elevational ranges approximate species realized niches, this conclusion conflicts with both the results presented here and a prominent verbal theory of how range limits are maintained in the absence of geographic barriers—that populations fail to adapt into novel niche space due to insufficient standing adaptive variation. This discrepancy may reflect selection bias in the species or genetic markers analyzed by Pennington et al., nonadditive effects on many functional phenotypes, or the influence of other phenomena like gene swamping. Teasing apart spurious patterns from causal mechanisms when autocorrelation among geography, climate, and morphology runs rampant remains a formidable challenge, but in systems that are unsuitable for experimental manipulation, we believe the careful scrutiny of patterns of variation in traits with demonstrated fitness consequences can be a powerful approach.

Macrophysiology is inherently a Faustian bargain of accepting noisy, imperfect data in exchange for the ability to reveal general phenomena that might otherwise remain hidden. This study is no exception, and we wish to highlight several factors that complicate any interpretation of our results. First, we would be remiss to conclude a paper on range limits without again emphasizing that they are multicausal phenomena: even if selection on hematological traits plays a major role in limiting elevational distributions, they are merely a handful stars in the larger constellation of interrelated physiological and anatomical traits that contribute to respiratory performance, a constellation itself positioned within the galaxy of other biotic and abiotic variables constraining jointly niche breadth. Second, variation in hematological trait values is itself influenced by numerous factors beyond elevation (Fair et al., 2007; Williamson & Witt, 2021), as elevational range dynamism and local adaptation shift optima and reaction norms. These factors, as well as unavoidable measurement error, are likely responsible for low effect sizes for predictors in our models, and correspondingly weak predictive power (**Table 1**). Integrating respiratory phenotypes with underlying genetic and evolutionary processes should therefore be a research priority in future studies investigating relationships between plasticity, adaptation, and niche breadth.

Nonetheless, we find the presence of correlations between physiology and biogeography at relatively fine scales a heartening step in answering one of biology’s most fundamental but challenging questions: why organisms live where they do.

## Acknowledgements

E.B.L. was supported by NSF DBI #1907353. Fieldwork in Peru was supported by National Science Foundation (DEB-1146491 and DEB-0543556) and permitted by SERFOR and its predecessor agencies. All research was conducted with UNM IACUC approval granted under protocols 16-200596-MC and 16-200418-MC. We thank Jon Nations for reviewing statistical methods and all code used in the study. For assistance in various aspects of amassing the dataset analyzed here, we thank Abraham Urbay T., Ashley Smiley, C. Gregory Schmitt, Christopher P. Barger, Daniel F. Lane, Donna C. Schmitt, Homan Castillo Benitez, Iris Olivas, Jennifer A. Clark, Jessica A. Castillo, Jimmy A. McGuire, Jose Ernesto, Huaroto Tornero, Luis Alza, Mariela Combe, Marlon Chagua, Matthew J. Baumann, Paloma Ordonez Buezo de Manzanedo, Robert Dudley, Robert J. Driver, Sabrina M. McNew, Selina M. Bauernfeind, Spencer C. Galen, Walter Vargas Campos, William A. Talbot and Zachary R. Hanna.

## Data accessibility

Specimen data are available from the Arctos database (https://arctosdb.org), associated with a forthcoming data paper (Witt et al., *in prep*.). All data and code used in this study are available on GitHub (https://github.com/elinck/andean_range_limits), along with a digital notebook documenting analyses.

